# NuFold: A Novel Tertiary RNA Structure Prediction Method Using Deep Learning with Flexible Nucleobase Center Representation

**DOI:** 10.1101/2023.09.20.558715

**Authors:** Yuki Kagaya, Zicong Zhang, Nabil Ibtehaz, Xiao Wang, Tsukasa Nakamura, David Huang, Daisuke Kihara

## Abstract

RNA is not only playing a core role in the central dogma as mRNA between DNA and protein, but also many non-coding RNAs have been discovered to have unique and diverse biological functions. As genome sequences become increasingly available and our knowledge of RNA sequences grows, the study of RNA’s structure and function has become more demanding. However, experimental determination of three-dimensional RNA structures is both costly and time-consuming, resulting in a substantial disparity between RNA sequence data and structural insights. In response to this challenge, we propose a novel computational approach that harnesses state-of-the-art deep learning architecture NuFold to accurately predict RNA tertiary structures. This approach aims to offer a cost-effective and efficient means of bridging the gap between RNA sequence information and structural comprehension. NuFold implements a nucleobase center representation, which allows it to reproduce all possible nucleotide conformations accurately.

## 1. Introduction

Ribonucleic acid (RNA) molecules are crucial for living organisms. One of the major roles of RNA is that of mRNA, one of the three components of the central dogma. Recent advances in high-throughput sequencing technologies and bioinformatics have unveiled a multitude of RNA molecules within cells that serve functions beyond mRNA ^1^. One of the most representative of these non-coding RNAs (ncRNA) are ribosomal RNA (rRNA) and transfer RNA (tRNA), which are important as part of a crucial machinery for translation from mRNA to protein. In addition to this, there are many varieties of ncRNAs that have their own unique functions, such as gene regulation or modification, and it is studied actively ^2,3^. Recently, these ncRNAs have been drawing attention, especially from the drug designing field by inhibiting or mimicking the activity of these functional RNAs ^4^. Knowledge of these ncRNA structures is critical to understanding the mechanism of function and designing brand-new RNA with specific functions since the three-dimensional structure directly affects its function. Understanding the three-dimensional structure of RNA is necessary for advancing RNA research in various ways. However, our knowledge of the RNA structures is limited compared to proteins, despite RNAcentral ^5^ keeping thirty million ncRNA sequences. For example, only around 6,000 entries include RNAs in the Protein Data Bank (PDB). This is only about 3 % of the entire PDB entries. Moreover, the Rfam database (v14.8) ^6^ has 4,094 families in their database, but only 124 (3.0 %) of them have at least one structure released in PDB. Therefore, computational methods predicting RNA tertiary structure with high accuracy have been demanded to close the gap between our sequence and structure knowledge.

Previously, a number of methods have been developed to predict RNA tertiary structure. There are two mainstream methodologies, template-based and simulation-based methods. Examples of template-based prediction methods include RNAbuilder ^7^ and ModeRNA ^8^. However, the three-dimensional structure of RNA known so far is even more limited compared to that of proteins, as we mentioned before, so this method has major limitations. Simulation-based prediction methods include SimRNA^9^ and FARFAR2 ^10^. These prediction methods sample while changing the structure to minimize internal energy. Although these methods are widely applicable compared to template-based methods, they require extensive computation to obtain good predictions. Also, the size should be less than a couple of hundred nucleotides to go through enough search space in a reasonable time. Furthermore, selecting the best structures from a large pool of thousands of structures remains a completely unresolved problem. Recently, ARES ^11^ has been proposed as a scoring method using deep learning that can solve this problem. Although ARES is not a structure prediction method per se, it can give a score to structures generated by other methods, increasing the likelihood of selecting a conformation that is better than the original energy score.

Here, we propose NuFold, our novel de novo RNA structure prediction method. NuFold takes an RNA sequence and produces a corresponding tertiary structure. Recently, AlphaFold^12^ marked a great performance in CASP14 (Critical Assessment of Structure Prediction) in 2020^13^, and this continues in CASP15 in 2022. CASP is the protein structure prediction competition held every two years, and more than a hundred participating groups will be given the sequence and submit its predicted structure. NuFold seeks to push the boundaries of current computational biology by making it possible to predict the three-dimensional structure of RNA from its sequence. NuFold implements nucleobase center representation to allow more flexible representation of each nucleobase. This allows us to reproduce any flexibility that exists in the base backbone, allowing us to create base shapes more accurately.

Three-dimensional structure prediction using machine learning is considered a difficult task because the available structural data is extremely limited compared to its spatial degree of freedom. In particular, the number of available RNA structures is even smaller than that of proteins, making this an even more difficult task. To overcome this difficulty, we employed a self-distillation technique to compensate for the lack of training data. This is a method that uses a network trained using existing data to assign labels to data that doesn’t have correct labels, and uses that data for further training of the network. It is known to be an effective method in the field of image recognition^14^. Here, we adopted a method that predicts the structure of an RNA sequence for which only the sequence is known, but the three-dimensional structure is unknown and utilized them as a true label in the training data.

## 2. Materials and Methods

### 2.1 Overview of the Nufold Architecture

Figure 1(a) shows the overall architecture of NuFold. NuFold shares the basic framework of deep learning architecture as AlphaFold, but with several key modification for folding RNA. In short, AlphaFold is an end-to-end protein structure prediction method from its sequence. AlphaFold takes a sequence, multiple sequence alignment (MSA), and template structures as the input features, and then directly outputs an atomic model of proteins. The AlphaFold framework consists of two major components, the Evoformer and the structure module. We can simply think of the former as an encoder and the latter as a decoder. In the first module, Evoformer, evolutionary information is extracted from the input MSAs and embedded into single and pair representations. Evoformer is trained in the same manner as the language model such as BERT, self-supervised by predicting masked words. Both representations are updated through Evoformer blocks by sort of attention networks. Pair representation especially keeps the information related to the relative position of each nucleotide. AlphaFold used 48 blocks of Evoformer. These two embeddings are used as input for the next module, the structure module, which consists mainly of the special attention mechanism called Invariant Point Attention (IPA). The structure module eventually predicts the rigid body translation and rotation of each base frame in Euclidean space along with updated single representation from both single and pair representation. AlphaFold used 8 blocks of the structure modules which shared the weights. AlphaFold is designed as end-to-end differentiable, losses came from every part of the network, including Evoformer’s BERT-style MSA loss, FAPE (Frame Aligned Point Error) loss which indicates structural misalignment, torsion angle prediction loss, and other auxiliary losses, are minimized during optimization.

**Figure 1.**
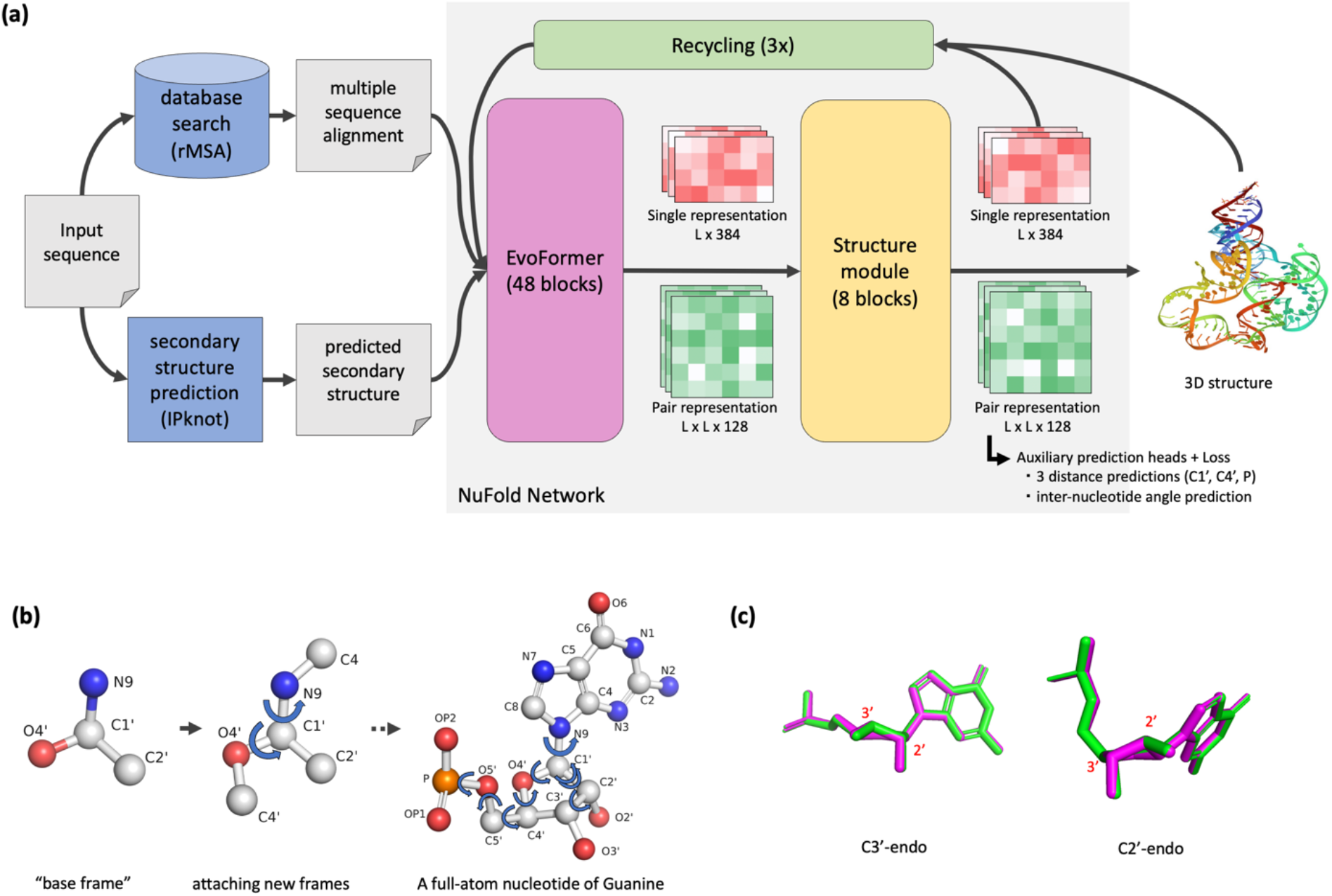
Overview of NuFold. (a) The architecture of NuFold. NuFold is an end-to-end architecture for RNA tertiary structure prediction, taking target sequence information and generating corresponding full-atom tertiary structures. The query sequence is initially used to construct a multiple sequence alignment (MSA), which, along with predicted secondary structure information, serves as input to the NuFold network. The NuFold network comprises two components: EvoFormer, a transformer model that extracts co-evolutional information from MSA and embeds it into both single and pair representations; and the structure module, which further processes the embedded information into 3D structures. These processes are iteratively performed in a recycling process to refine predictions. (b) The network predicts two key components for full-atom structure prediction: the translation and rotation of the base frame, along with a set of torsion angles derived from the base frame. These torsion angles are used to extend new atoms. (c) Assessing sugar puckering reproducibility. Green and magenta structures represent ground truth and predicted structures, respectively. NuFold predicts torsion angles not only for the main chain or chi angles but also for the ribose ring, allowing for the reproduction of sugar-puckering formations.

To be able to predict RNA tertiary structures, we implemented several key modifications atop the original AlphaFold framework. One of the most significant alterations pertains to the reference definition of RNA nucleotide shapes (A, C, G, and U), including considerations for the relative distances between atoms and hinge angles. In the NuFold approach, we define the base frame using four atoms: O4’, C1’, C2’, and the first N of the base (N1 for C and U, N9 for G and A). All other atoms are partitioned into ten frames, which are then iteratively bonded using predicted torsion angles on the bonds between frames as a guiding principle (as illustrated in Figure 1b). It’s worth noting that these definitions were initially hardcoded for the 20 amino acids in AlphaFold and needed adaptation for RNA. This representation of the RNA nucleotides with frames and torsion angles can reproduce the full dynamics of nucleotide shapes (see Figure 1c). Moreover, we made changes in the character types (words) used for Multiple Sequence Alignment (MSA) within the Evoformer. The character type count transitioned from 20 + 3 (standard amino acids + unknown + gap + masked) to 4 + 3. We also introduced new auxiliary heads that consider the distance between C4’ and P, as well as the dihedral angle between residue pairs. These auxiliary networks were constructed using three layers of 2D convolution, departing from the original linear layer used in AlphaFold. The input for NuFold is the sequence of query, MSA, and predicted secondary structure produced by IPknot ^15,16^ and the in-house modified version of IPknot, and the output is the all-atom coordinates of its structure. AlphaFold was using information from templates, but it is substituted by secondary structure in NuFold. Given the limited availability of templates for RNA, we removed the template input network and adapted it to accept the predicted secondary structure as input. Additionally, we adopted C1’ carbon for tasks originally associated with Cα, such as lDDT-Cα, and N1/N9 for tasks initially related to Cβ, such as the original distogram head. The NuFold network consists of two major components, the Evoformer and the structure module with the same number of blocks. Loss is also similar to AlphaFold, but some items have been added and the weight has been changed to accommodate NuFold’s unique extensions (see section 2.4). In addition, NuFold is adopting a well-engineered system from AlphaFold, such as recycling, which reuses the final output as input to iteratively improve its prediction.

Moreover, we utilized metagenome sequences for MSA construction. The quality of the sequences in the metagenome sequences is not always that good, but it is known that using those sequences to accumulate depth and diversity in the MSA increases the performance of the structure prediction. We gathered metagenome sequences from public datasets to construct our own metagenome sequence database for MSA construction.

### 2.2 Dataset Construction

For the training dataset, we sourced RNA tertiary structures from the Protein Data Bank (PDB) ^17^. Our selection criteria included structures solved through X-ray crystallography or Cryo-electron microscopy with a resolution better than 5.0 Å and a sequence length ranging from 10 to 1500 nucleotides. Additionally, we applied a masking procedure to exclude bases that were in close spatial proximity to other RNAs or proteins, retaining only strands with 10 or more consecutive unmasked bases. To ensure structural stability, we required each RNA to contain one or more standard Watson-Crick base pairs in non-masked regions. After applying these criteria, we identified 3,237 RNA chains that met our requirements. We supplemented this dataset with RNA chains used in the RNA-puzzles competition ^18^ and performed sequence clustering. Clustering employed complete linkage, considering global sequence identity between different RNA chains. Chains with a sequence identity exceeding 80% were grouped together. We used single-linkage clustering to ensure that distinct clusters did not contain similar RNA chains. This process resulted in a total of 499 clusters. From these clusters, 403, 48, and 48 clusters were allocated for training, validation, and testing, respectively. We also excluded targets from the test set that were not RNA-puzzles monomers, resulting in a final test set size of 36. The training dataset was expanded to encompass all the structures from 2,860 chains.

We employed the bpRNA-1m ^19^ dataset to construct a semi-supervised training dataset following the principles of Noisy Student learning. The need for a larger dataset was driven by the desire to enhance generalization performance, given the relatively small number of examples in the clean PDB dataset. Similar to our approach with the PDB dataset, we conducted sequence clustering on the bpRNA-1m dataset and selected 11,197 sequences distinct from those in the PDB dataset. Subsequently, we used trRosettaRNA ^20^ to predict three-dimensional structures for these sequences, focusing on those with one or more Watson-Crick base pairs. This process yielded a total of 11,101 structures, which served as our self-distillation dataset.

### 2.3 Input Feature Generation

We utilized two input features for our training. The first input feature was generated using multiple sequence alignments (MSA), which served as a source of co-evolutional information. To create the MSAs, we employed the rMSA pipeline ^21^ with default parameters. This pipeline leverages data from the RNA central database, NCBI nt database ^22^, and Rfam database ^6^, utilizing BLAST ^23^ and infernal algorithms ^24^ for searching. The second input feature incorporated predicted secondary structures. We employed IPknot to generate these predictions. To account for the lack of confidence or significance information in IPknot’s output, we introduced a stochastic process. In this modified IPknot version, we randomly discarded 10% of the constraints and optimized the objective function through integer programming. This process was repeated ten times to identify the base pairings essential for valid RNA structures. The resulting binary base-pairing matrices were averaged to estimate the likelihood of base pairing between specific bases. This resulting LxL matrix, where L represents the length of the RNA chain, also served as an input feature for the predicted secondary structure.

### 2.4 Training of NuFold

In the training of NuFold, we followed a methodology similar to that of AlphaFold, with some minor distinctions. Our loss function is defined as:

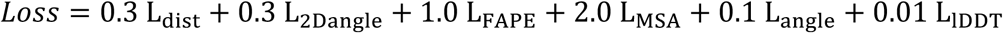

This loss function comprises several components. *L* _*dist*_ represents the aggregation of losses from three distogram heads, aiming to predict 40 evenly spaced distance bins spanning from 2 Å to 22 Å, and loss was computed using cross-entropy. *L*_2*Dangle*_ represents the loss from an auxiliary head responsible for predicting torsion angles between pseudo-bonds of two bases. This loss is newly introduced in this study, and this involves predicting 24 even-sized bins with cross-entropy loss. *L*_*FAPE*_, *L*_*MSA*_, *L*_*angle*_, and *L*_*lDDt*_ encompass the FAPE loss, BERT-like MSA loss, torsion angle loss, and plDDT prediction loss, respectively. We retained these components without modification.

During the fine-tuning step, we introduced a structural violation loss, denoted as *L*_*viol*_, which penalizes atom clashes and longer bonds. The total loss during fine-tuning becomes:

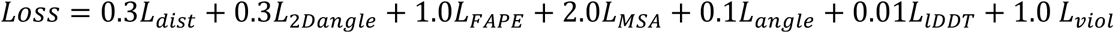

For the training process, we utilized the Adam optimizer with a learning rate of 1e-3 for the initial training phase and 1e-4 for fine-tuning. The training was conducted in two steps. First, we trained without the violation loss. Once the loss converged, we proceeded with fine-tuning by increasing the weight of the violation loss. Unlike AlphaFold, our training did not incorporate a warm-up step. The batch size was set at 16, with gradient accumulation and gradient clipping applied at 0.01.

### 2.5 Evaluation Metrics

In our evaluation, we relied on several key metrics to assess the accuracy of our predictions. C1’-RMSD (Root Mean Square Deviation) is a widely used metric that quantifies the structural similarity between the predicted and ground truth structures. It measures the average deviation between corresponding atoms in the two structures. GDT-TS (Global Distance Test) is another metric used to evaluate the accuracy of the global structure. It measures the overall structural similarity between the predicted and ground truth structures. lDDT (local Distance Difference Test) assesses the accuracy of the local structure. It provides insights into how well the predicted local structure aligns with the ground truth.

In addition to these metrics, we employed the Interaction Network Fidelity (INF) metric, as used in RNA-puzzles assessment ^25^. INF considers the entire RNA structure as an interaction network consisting of WC interactions, non-WC interactions, and base stacking. INF is defined as the Matthews correlation coefficient (MCC) of this network between the reference and predicted structures. A high INF score means that the relative positions of the bases are correctly reproduced.

## 3. Results

### 3.1 Analysis of Structure Prediction Performance

First, we selected epochs with the best and second-best averaged RMSD in the validation dataset, which is named “ave. RMSD #1” and “ave. RMSD #2” in Table 1, respectively. Also, we selected the best GDT-TS performance, which is named “ave. GDT-TS #1” in Table 1. Finally, we selected the epoch based on the best Rosetta score, which is named “ave. Rosetta #1” in Table 1 Additionally, we implemented a structural clustering strategy for the structures generated during different training epochs. The clustering aimed to mitigate the influence of outliers and reduce the possibility of selecting suboptimal structures. Two specific strategies emerged from this clustering. The first strategy selects the structure within the largest cluster that is closest to the cluster’s average structure, which is named as “cluster #1” in Table 1. The second strategy chooses the structure with the lowest Rosetta score from the largest cluster, which is named as “cluster #2” in Table 1. We also considered a straightforward strategy that employs plDDT scores to select structures from a pool of candidates, which is named as “plDDT” in Table 1. While a larger structure pool may contain better structures, it also demands more computational resources to expand the pool size. Table 1 reports C1’-RMSD and GDT-TS metrics for the test datasets. Notably, the #2 RMSD model exhibits better overall performance in terms of both RMSD and GDT-TS. Consequently, this model serves as the baseline for subsequent analyses.

**Table 1.** Overall performance of NuFold using different model selection criteria.

| Criteria | Steps | RMSD (Å)<br>(validation) | GDT-TS<br>(validation) | RMSD (Å)<br>(test) | GDT-TS<br>(test) | # better than<br>5 Å (test) | # win to<br>baseline<br>(win / tie /<br>lose) |
| --- | --- | --- | --- | --- | --- | --- | --- |
| ave. RMSD #1 | 124863 | 6.38 | 0.593 | 7.66 | 0.437 | 19 / 36 | 4 / 26 / 6 |
| ave. RMSD #2 | 146287 | 6.43 | 0.598 | 7.68 | 0.442 | 20 / 36 | Baseline |
| ave. GDT-TS #1 | 145263 | 6.48 | 0.598 | 7.55 | 0.443 | 19 / 36 | 5 / 29 / 2 |
| ave. Rosetta #1 | 110095 | 6.92 | 0.582 | 7.98 | 0.430 | 19 / 36 | 7 / 22 / 7 |
| cluster #1 | - | 6.56 | 0.467 | 7.81 | 0.433 | 19 / 36 | 5 / 26 / 5 |
| cluster #2 | - | 6.57 | 0.466 | 7.80 | 0.434 | 19 / 36 | 5 / 26 / 5 |
| pLDDT | - | 6.51 | 0.471 | 7.79 | 0.451 | 22 / 36 | 7 / 24 / 5 |

The criteria “ave. RMSD #1” and “ave. RMSD #2” is the best and second best model selected by average RMSD performance using validation dataset. “ave. GDT-TS #1” is the best model selected by the average GDT-TS score using validation dataset. “ave. Rosetta #1” is the best model selected by the average Rosetta score using validation dataset. “cluster #1” is the strategy that first structures generated by each network weights were clustered, and picked up the structure from the largest cluster which is closest to the averaged structure in the cluster. “cluster #2” is the same clustering strategy as previous, but the lowest Rosetta score structure was selected from largest cluster. “pLDDT” is the strategy which we selected the structure which has highest pLDDT score from the structures.

To illustrate the characteristics of the structure produced by NuFold, we investigated the relationship between performance and factors that can directly affect prediction performance. Figure 2 shows the relationship between RMSD and (a) length of RNA, (b) number of sequences in MSA, and (c) effective sequence count in MSA, respectively. These analyses provide valuable insights into how specific target characteristics correlate with the accuracy of RNA structure predictions. We observe that longer RNA chains tend to give relatively larger RMSDs, but this relationship is not clear (Figure 2a). Larger MSAs provide more evolutionary insight and should therefore be inherently more prone to accurate predictions than smaller MSAs, but this trend was also not clearly observed (Figure 2b and c).

**Figure 2.**
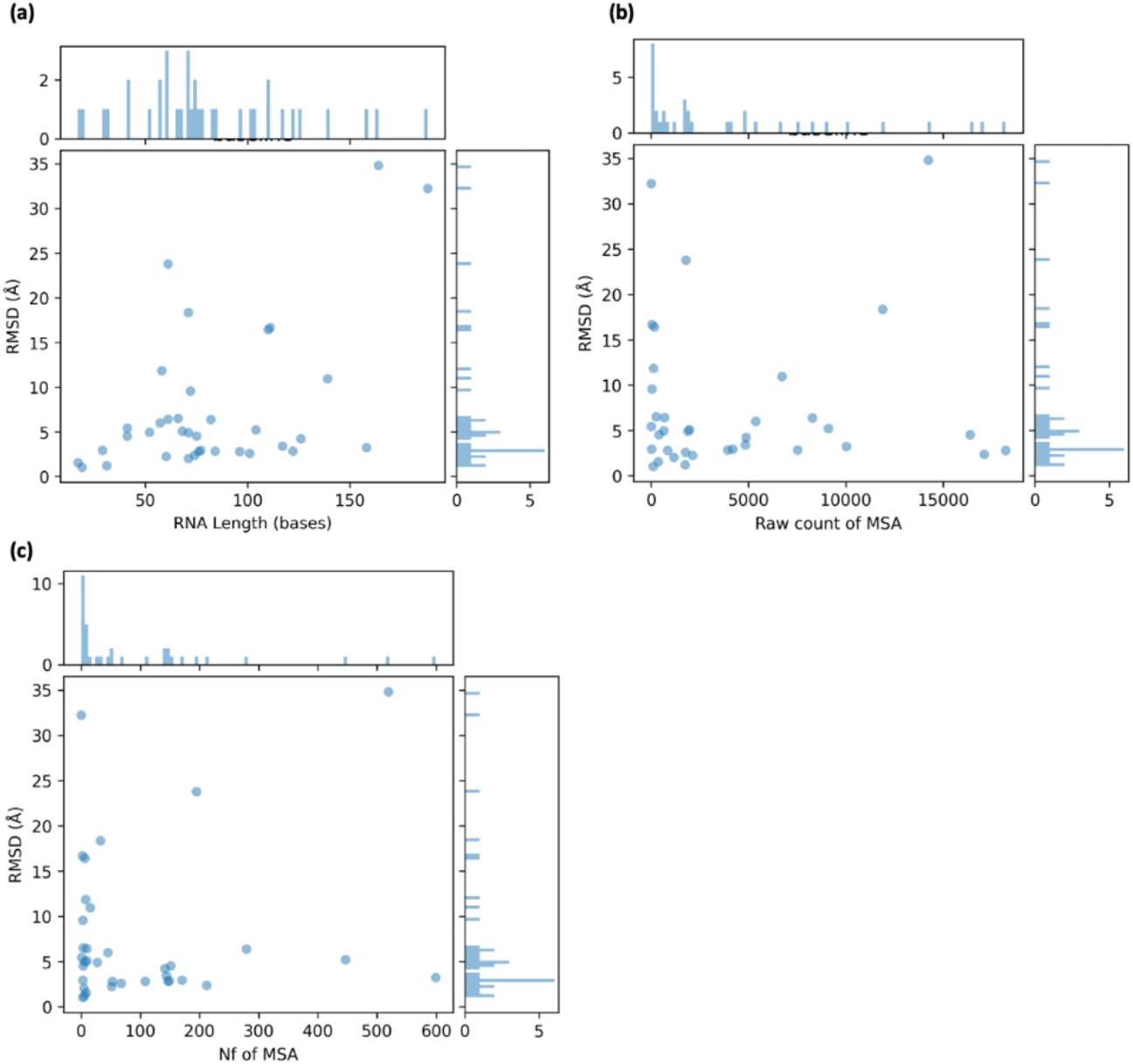
Comparison between RMSD and characteristics of targets. Lower RMSD means more accurate structure prediction. (a) Comparison between the length of query RNA sequence and RMSD. (b) Comparison between the depth of MSA and RMSD (Å). (c) Comparison between the number of effective sequences (Nf) of the MSA and RMSD.

### 3.2 Effect of Recycling

NuFold implements the same recycling mechanism as AlphaFold. After the first prediction is made, recycle mixes the output atomic coordinates and internal representation vectors with the original input and uses them as input for the network. This technique can be applied repeatedly, and the network is expected to improve its predictive performance by using its output as a guide. During training, 0 to 3 recycles were randomly selected for each batch. In inference, as a default, he uses three recycles. Although only up to three recycles were used during training, previous work revealed that increasing the number of recycles improves prediction performance for many proteins when analyzing Alphafold results 26. To investigate the effect of recycling on NuFold, we performed 30 recycles and examined the predicted performance at each recycle number. The relationship between the number of recycles and RMSD and plDDT is shown in Figure 3 (a). The performance gradually improves along with the number of recycles up to 17, but it ends up going worse with more number of recycles. Figures 3 (b), (c), and (d) show the comparison between the baseline, 17 times recycling, and 30 times recycling and selecting max plDDT model each other. Although the optimal number of recycles differs depending on the query, we can see that it is possible to select the optimal structure using plDDT after performing enough recycles.

**Figure 3.**
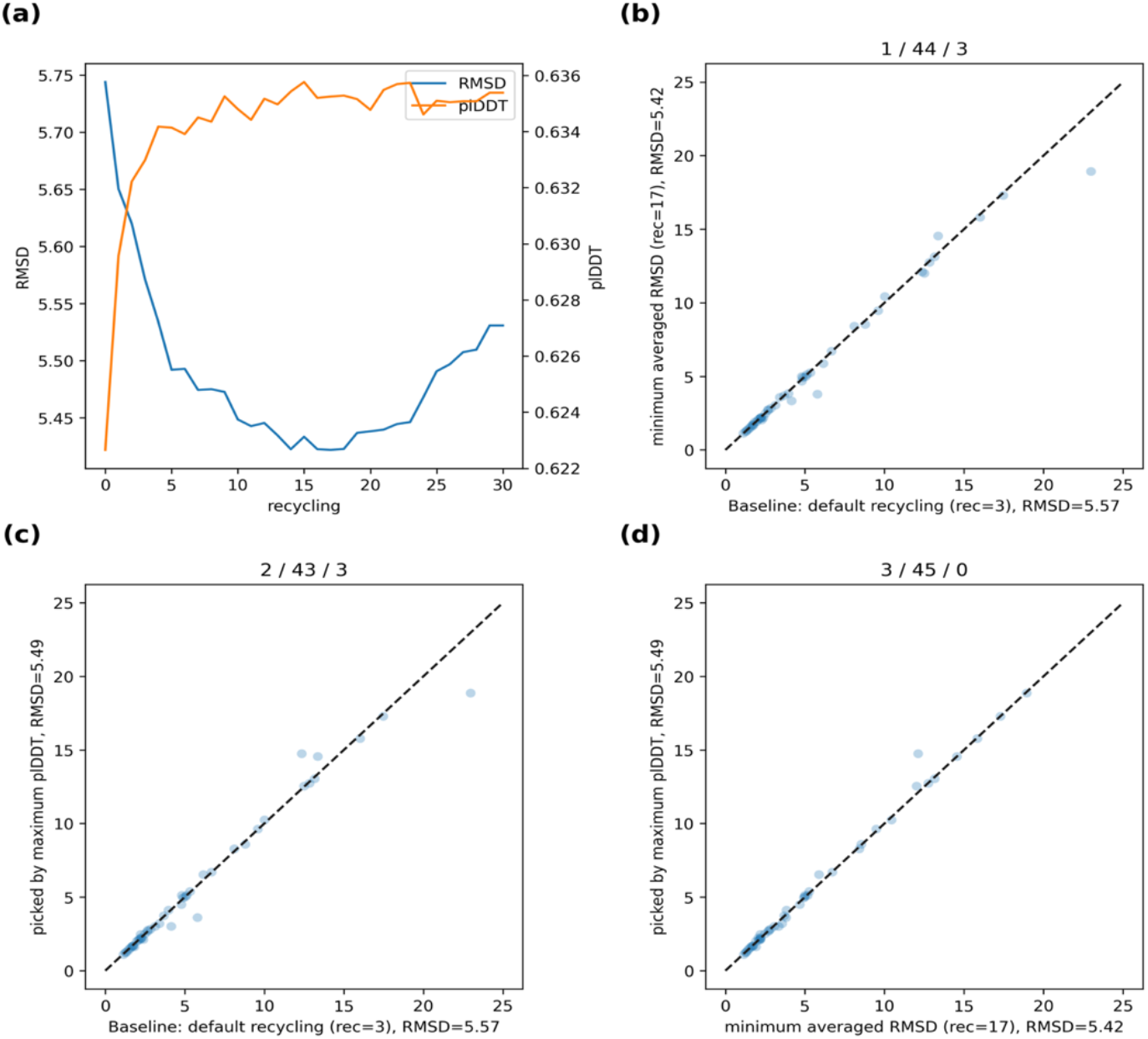
The effect of recycling. (a) The relationship between the number of recycling against RMSD (left axis, Å) and plDDT (right axis). (b) Comparison between the default recycling (rec=3, baseline) and minimum averaged recycling (n=17). The unit for both the x and y axes is RMSD in Å. The three numbers in the title of each plot correspond to the number of points above the diagonal line, the number of points on the line, and the number of points below the diagonal line, respectively. (c) Comparison between the default recycling (rec=3, baseline) and the models picked by plDDT from all 30 recycling. (d) Comparison between minimum averaged recycling (n=17) and the models picked by plDDT from all 30 recycling.

### 3.3 Effect of Using Metagenome Sequence for MSA

At Alphafold, MSA depth is known to be a crucial factor for its performance. Since EvoFormer is designed to extract co-evolutionary information between residues that is useful for structure prediction from MSA, prediction performance is generally greatly reduced when MSA has little information. Since NuFold has a very similar architecture and utilizes MSA as its primary source of information, we can reasonably infer that MSA depth affects its predictive performance.

Since NuFold utilizes rMSA as MSA, its sequence database is limited to Rfam, RNA central, and NCBI nt. One breakthrough in the field of protein structure prediction is the observation that the use of metagenomic sequences improves prediction performance. Although NCBI nt partially contains metagenomic sequences, we decided to extend the sequence database to allow more metagenomic sequences to be available as MSAs.

As a metagenome database, in addition to NCBI env_nt, we created a database using TARA Ocean Metagenome ^27^, Mgnify MAG ^28^, and all Mgnify contigs. The total size is around 3.0 Tb, but this includes redundancy. Database searching is achieved by further extending the rMSA pipeline, similar to the last stage of rMSA, but essentially similar to the RNAcmap pipeline^29^. Briefly, the query sequence first searches the metagenome sequence database with BLASTn, and then creates a covariance model (CM) with infernal from the results. This CM is used as input for infernal’s cmsearch, which searches the same metagenomic database. Finally, this resulting MSA is combined with the rMSA result to remove redundancies with hhfilter.

Overall, we showed that prediction performance can be improved by mixing metagenomic sequences with the original MSA. However, no correlation was found between increasing MSA size and increasing accuracy. Although the size of the MSA could not be an indicator to predict the structure prediction performance in advance, the plDDT of the prediction result was found to be an indicator that can be used to better select an MSA. Figure 4 shows a comparison of the baseline performance and the performance when selecting the largest plDDT from among the predicted structures of each of the four MSAs. By selecting the structure that maximizes plDDT from four inferences, the average RMSD improves by 1.05 Å. Also, we could achieve the same or better RMSD with 28 targets (78%), but with 2 targets, the accuracy is lower with more than 0.5 Å margin. Overall, it can be seen that the metagenome sequence has the effect of improving structure prediction performance. What is noteworthy here is that the prediction performance is improved even though no metagenomic MSA is used during training. By using metagenomics during learning and retraining, metagenomic interpretation performance may improve and further performance improvements may be seen.

**Figure 4.**
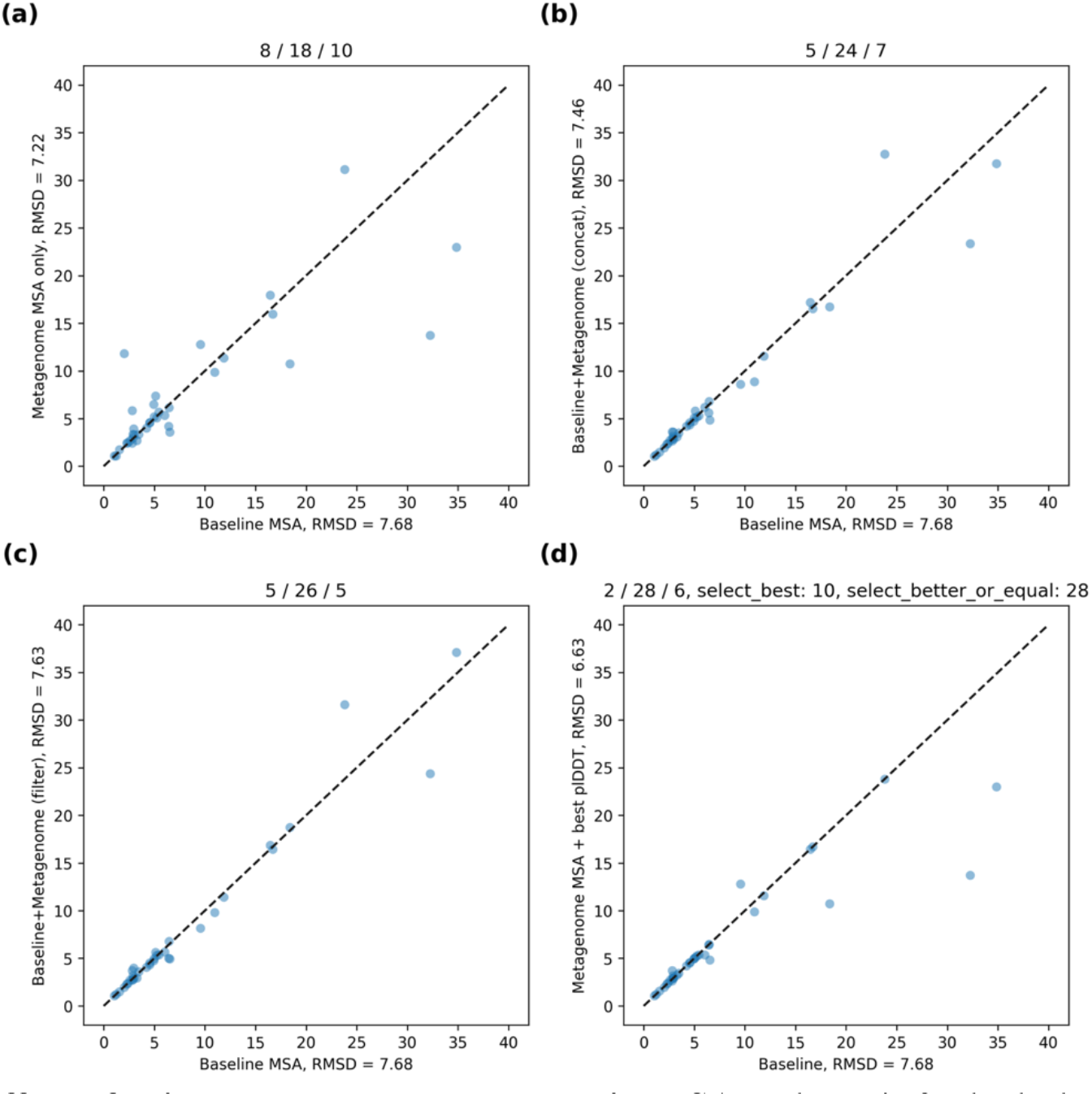
The effect of using metagenome sequences in MSAs. The unit for both the x and y axes is RMSD in Å. The three numbers in the title of each plot correspond to the number of points above the diagonal line, the number of points on the line, and the number of points below the diagonal line, respectively. (a) Comparison between the default MSA (baseline) and inference using only metagenome MSA. (b) Comparison between the default MSA (baseline) and inference using metagenome MSA concatenated with original MSA. (c) Comparison between the default MSA (baseline) and inference using metagenome MSA filtered after concatenated with original MSA. (d) Comparison between the default MSA (baseline) and models picked by plDDT from 4 different MSAs including original MSA.

### 3.4 Case Studies

In order to better understand NuFold’s performance, we present concrete examples illustrated in Figure 5. The first example is rp06 from RNA-puzzles (Figure 5a): This target is a 168-base adenosylcobalamin riboswitch with PDB ID: 4GXY. NuFold achieved an all-atom RMSD of 3.4 Å. Notably, as of RNA-puzzles round 2 ^30^, this target had a top all-atom RMSD of 11.4 Å, marking a significant advancement by NuFold. The challenge with this relatively large riboswitch lay in efficiently determining the correct structure using traditional methods. Evaluation of RNA-puzzles indicated that the absence of a ligand to fill the riboswitch pocket could be the cause of incorrect predictions, although this effect was rarely observed in NuFold prediction.

**Figure 5.**
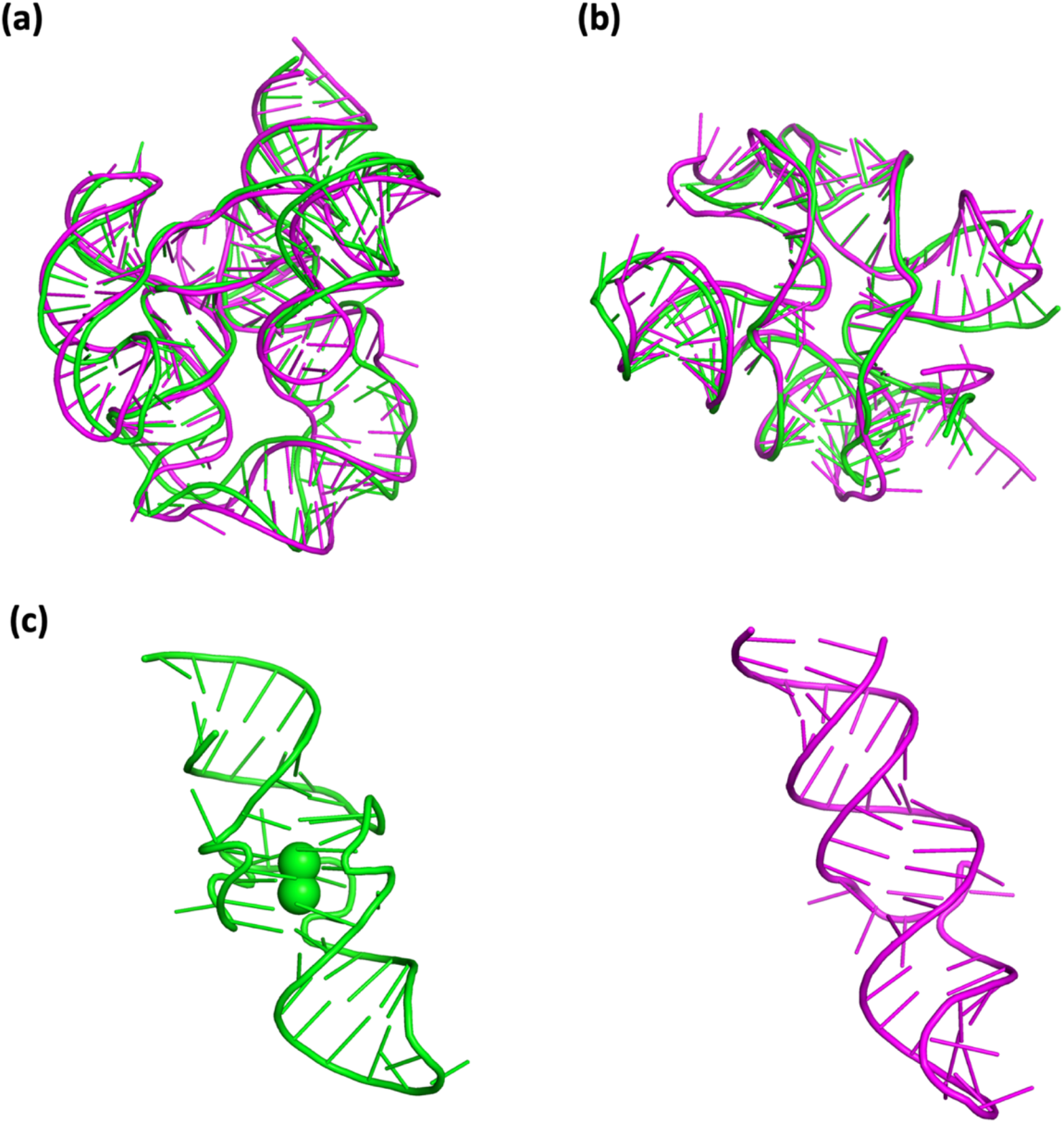
Case studies of the predictions of NuFold. The native structures and predicted structures are superimposed and shown in green and magenta, respectively. (a) The prediction of rp05. (b) rp12 (c) One of the failed cases with G-quadruplex local structure. The native structure contains two potassium ions in the middle of two stories G-quadruplex motif. NuFold prediction (right, magenta) could not reproduce this motif and got 12 Å RMSD.

The second example is rp12 from RNA-puzzles (Figure 5b): This target is a 108-base ydaO riboswitch with PDB ID: 4QLM. While the structure of a c-di-GMP-binding riboswitch with a similar function had been previously solved at that time, the ydaO riboswitch presented a novel structural topology that made this target challenging to predict. NuFold successfully predicted this structure with an all-atom RMSD of 3.5 Å. Remarkably, as of RNA-puzzles round 3 ^31^, the top all-atom RMSD for this target was 10 Å, indicating a significant improvement in performance by NuFold. Changes in available structural data may alter the target’s difficulty over time, but it’s worth noting that previously challenging targets can now be predicted with practical accuracy.

The third example is 7OA3 from PDB. This case serves as a notable failure included in the validation dataset. The target is a relatively small RNA aptamer consisting of 52 bases, notable for its local G-quadruplex structure at the center. NuFold obtained an all-atom RMSD of 9.1 Å. NuFold tends to predict general RNA double helix structures, and accurately predicting G-quadruplex structures remains a challenge due to their distinctiveness. Given the limited training data, generalizing such rare structures can be challenging. Such local structures must be able to be accurately predicted, as they may be essential for elucidating their function.

## 4. Discussion

In this paper, we have introduced NuFold, a novel de novo structure prediction method for RNA. The three-dimensional structures of non-coding RNAs (ncRNAs) are intricately linked to their functions, mirroring the relationship between structure and function in proteins. However, unlike proteins, the available experimental structures for ncRNAs are more limited in variety and quantity. This disparity has underscored the pressing need for accurate and computationally efficient methods for predicting RNA structures. NuFold draws inspiration from the architectural success of AlphaFold, a renowned protein structure prediction method, and shares the basic computational framework. Yet, the development of machine learning-based RNA structure prediction methods faces a unique challenge: a paucity of structural data for training. To surmount this challenge, we leveraged self-distillation data, a strategy akin to NoisyStudent, to augment our training dataset and enhance the predictive power of NuFold. Additionally, we were able to confirm that NuFold’s recycling function was effective. It was observed that having a sufficient number of recycles is important for better performance, but it was also observed that too many recycles had a negative impact on performance. We also observed that enriching MSA with metagenomic sequences contributed to significant performance improvements. For both strategies, plDDT was found to be an efficient metric for selecting good predictive structures.

While the results achieved by NuFold may not reach the same groundbreaking level as those of AlphaFold for proteins, several noteworthy distinctions emerge from our observations. Notably, we observed variations in certain aspects, such as the relationship between Multiple Sequence Alignment (MSA) depth and predictive performance. Unlike AlphaFold, where MSA depth has been reported to correlate with performance, NuFold did not exhibit a significant relationship in this regard. Several factors may contribute to this disparity.

One possible explanation lies in the intrinsic differences between RNA and protein structure prediction. For RNA secondary structure prediction, which is similar to contact prediction in the protein field, there exists a baseline level of predictive performance even in methods that do not rely heavily on MSA. This contrasts with protein structure prediction, where MSA depth plays a more critical role. RNA secondary structure prediction benefits from the inherent nature of RNA, where the formation of hydrogen bonds between base pairs largely governs thermal structural stability. This problem can be addressed with dynamic programming techniques traditionally, offering a more straightforward solution compared to the complexities involved in protein folding.

Additionally, the diverse structural motifs found in RNA molecules, including bulges, internal loops, and pseudoknots, present unique challenges. These motifs often deviate from the regular secondary structure patterns commonly seen in proteins. Therefore, NuFold’s architecture and training approach can be tailored to account for these distinctive RNA structural features more. Furthermore, the varying degrees of structural conservation across RNA families can influence the effectiveness of MSA in predicting RNA structures. Unlike proteins, where conserved structural motifs are more prevalent, RNA structures can exhibit greater variability even within closely related sequences, making it challenging to extract universally applicable rules from MSAs.

In recent bioinformatics, the application of machine learning has paved the way for solving a multitude of problems. This progress hinges on a symbiotic relationship between the extensive data collected by experimentalists and the continuous development of machine learning techniques. While AlphaFold’s architecture stands as a testament to engineering ingenuity in the realm of protein structure prediction, the limited availability of RNA data underscores the necessity for a multifaceted approach. To propel RNA structure prediction forward, the field demands not only increased data but also innovative architectural advances. These dual imperatives represent the path toward further breakthroughs in the field of computational biology.

## Acknowledgements

This work was partly supported by the National Institutes of Health (R01GM133840, R01GM123055, and 3R01GM133840-02S1) and the National Science Foundation (CMMI1825941, MCB1925643, DBI2003635, and DBI2146026) to DK.

## Author contributions

DK conceived the study. ZZ has modified the distributed Alphafold2 code and made it trainable. YK designed and implemented NuFold based on the trainable Alphafold2 code developed by ZZ. NI modified the source of ipknot to output fraction values for nucleotide base pairs. XY implemented the refinement pipeline. TN performed benchmarks. YK performed the computation and YK and DK analyzed the data. YK drafted the manuscript and DK edited it. All the authors read and approved on the manuscript.

## Data and Code Availability

The trained model we used in the paper and all the codes for inference will be available after we clean up the codes.

